# Is YouTube^®^ promoting the exotic pet trade? Analysis of the global public perception of popular YouTube^®^ videos featuring threatened exotic animals

**DOI:** 10.1101/2020.06.17.156661

**Authors:** Georgia Moloney, Jonathan Tuke, Eleonora Dal Grande, Torben Nielsen, Anne-Lise Chaber

## Abstract

The exploitation of threatened exotic species via social media challenges efforts to regulate the exotic pet trade and consequently threatens species conservation. To investigate how such content is perceived by the global community, sentiment analysis techniques were employed to explore variations in attitudes expressed through text and emoji usage in public comments associated with 346 popular YouTube^®^ videos starring exotic wild cats or primates in captive or domestic situations. Although a negative trend in mean text sentiment was observed in 2015 for primates, an otherwise consistent positive mean sentiment text and emoji score through time was revealed in response to both exotic wild cat and primate videos, implying the societal normalisation and acceptance of exotic pets. These findings highlight the urgency for effective YouTube^®^ policy changes and content management to promote public education and conservation awareness, whilst extinguishing false legitimisation and potential demand for the exotic pet trade.

## Introduction

Unsustainable trade in wildlife has been recognised as an important challenge to maintaining species conservation, where live animals are trafficked for pets amongst other purposes [1]. Exotic pets have been defined as animals without an extensive history of domestication or life in captivity that are not traditionally viewed as companion animals [2]. The exotic pet trade encompasses the global exploitation of exotic animals often sourced from wild populations, thus imposing a significant threat to species biodiversity, animal welfare and public health [2–4]. Although international wildlife trade regulations are in place through the Convention on International Trade in Endangered Species of Wild Fauna and Flora (CITES), much of the trade still operates illegally [1,3,5].

Social media has been identified as an important driver of the trade through influencing public perception [2]. Social media has altered how society access, consume and share information, wherein exposure to content is individually tailored to enhance user experience [6]. Such platforms have become a dominant source for news, entertainment and information, therefore understanding factors which generate appeal for these sites is important to enable content regulation [7,8]. Currently, a considerable amount of published content remains unregulated due to poorly established policy guidelines and limited enforcement disproportionate to the volume of data uploaded [9].

With more than two billion users worldwide as of 2020, YouTube^®^ is the largest global online video website and is thus considered to be one of the most influential social networking platforms [10,11]. YouTube^®^ enables users to upload, create, share and engage in content, thus promoting global participation [6]. Individuals accessing insufficiently regulated videos on YouTube^®^ commonly choose to express their opinions and approval or condemnation of content through written language and emoji usage within the associated comments section [12]. Therefore, these comments may indicate how the public perceives content depicting exotic animals in a variety of captive or domestic situations and further suggest how this perception is altered throughout the life of a video. Sentiment analysis, being the study of opinions, attitudes and emotions conveyed via text, may be employed to evaluate positive or negative intentions associated with user comments [7,13,14]. It has been suggested that YouTube^®^ may be used to influence public perception surrounding ‘cute’ exotic animals and thus promote demand for exotic pets, however the significance of this platform as a driver of the exotic pet trade has not been fully investigated [15].

The aims of this study were to employ sentiment analysis techniques to explore public perception of exotic species featured in popular YouTube^®^ videos, variation in the perception throughout the life of a video and examine the potential association with the exotic pet trade and impact on species conservation. Two of the most commonly exploited mammalian orders include primates and carnivores (including Felidae) [2], hence these groups were targeted as the basis for examining videos for analysis. YouTube^®^ policies encompassing animal usage will be assessed accordingly.

## Materials and methods

### Online Data Collection

Common wild cat and primate species kept as exotic pets were manually researched, predominately through Google^®^ and with reference to the International Union for Conservation of Nature (IUCN) Red List [16], and utilised as search terms within YouTube^®^. Popular terms which formed the basis of the video search included: ‘cheetah’, ‘tiger’, ‘lion’, ‘leopard’, ‘jaguar’, ‘monkey’, ‘capuchin monkey’, ‘marmoset monkey’, ‘primate’, ‘chimpanzee’, ‘slow loris’, ‘spider monkey’, ‘macaque’, ‘lemur’ and ‘gibbon’. Further variations in search terms included the addition of seemingly popular key phrases, including ‘pet’, ‘cute’ and ‘baby’. A full list of search terms investigated is located within the supporting information (S1 Table).

Inclusion criteria involved the portrayal of an exotic animal either within a captive (i.e. zoo) or domestic (i.e. pet) setting, interacting in some manner with either a human or another species. Videos were only considered if they had accumulated a minimum of one million views at the time of collection to ensure they had been received by a significant audience. Videos were excluded from the study if the associated comments section had been disabled. Videos published between May 2006 and October 2019 were included within the study.

Once selected, the following information was manually extracted and entered into a Microsoft Excel spreadsheet: Video web address (URL), search term used, species identified, IUCN Red List status, CITES Appendices listings, setting (captive vs pet) and species interactions present. Species were determined through analysis of the animal(s) featured in comparison to images and information sourced through online research (via Google^®^) and the IUCN Red List, unless indicated within the video title or description.

Data management and analysis was conducted using R Studio (version 3.6.1) [17]. Additional data was extracted through the *tuber* package [18] to fulfil the following categories: Video ID, comments, date published, date downloaded, view count, like count, dislike count and comment count. The data was arranged to ensure each row represented an individual comment for analysis using the *tidytext* package, accompanied by the language detected through the *textcat* package [19,20]. Video information and comments were extracted from May 2006 to October 2019. In accordance with the aims of the study, the comments produced over time for each video were evaluated for sentiment based on language (English only) and emoji usage.

A multiple-response approach was applied to analyse species frequencies, taking into consideration the number of instances in which a particular species was featured across all videos within their assigned animal group. For example, when selecting for videos using primate specific search terms, some videos included multiple primate species, where each primate species appearance was listed as an observation. Twelve primate videos were considered to be ‘compilation videos’ as they featured four or more videos compiled together. In these instances, species and IUCN Red List classification were not specified due to ambiguity.

### Text sentiment Analysis

Sentiment analysis techniques were employed to overcome limitations imposed by traditional manual qualitative techniques and enable more efficient analysis of a large dataset [21]. Lexicons derived from the *tidytext* package were applied to impart values to key words [20,22]. The “afinn” lexicon provided negative or positive values between −1 and +1 to particular pre-determined sentiment terms to quantitatively analyse variation in mean text sentiment across comments. Similarly, the “bing” lexicon qualitatively analysed phrases, denoting words as either ‘negative’ or ‘positive’, and was thus used to analyse word frequencies. Due to software limitations, only English comments were analysed for text sentiment. Trends in text sentiment were compared with emoji sentiment.

### Emoji sentiment analysis

Comments containing emojis were extracted and analysed within R Studio (version 3.6.1) [17]. Due to the over-representation of wild cat comments containing emojis, a random sample of 10,000 comments were selected (Fig 1). The emoji sentiment lexicon utilised in this paper was based on work conducted by Novak et al. [14] and Peterka-Bonetta [23]. The lexicon provided an average sentiment score for each emoji-containing comment as a value between −1 and +1, wherein negative values represent negative emoji usage and positive values correspond to positive emojis. Changes in sentiment associated with emoji usage over time (based on comment publication dates) were analysed in relation to species and conservation status. Filters were applied to ensure comments analysed contained at least one emoji and a sentiment value not equal to zero. Mean emoji sentiment scores through time were further analysed in conjunction with mean text sentiment scores.

**Fig 1.**
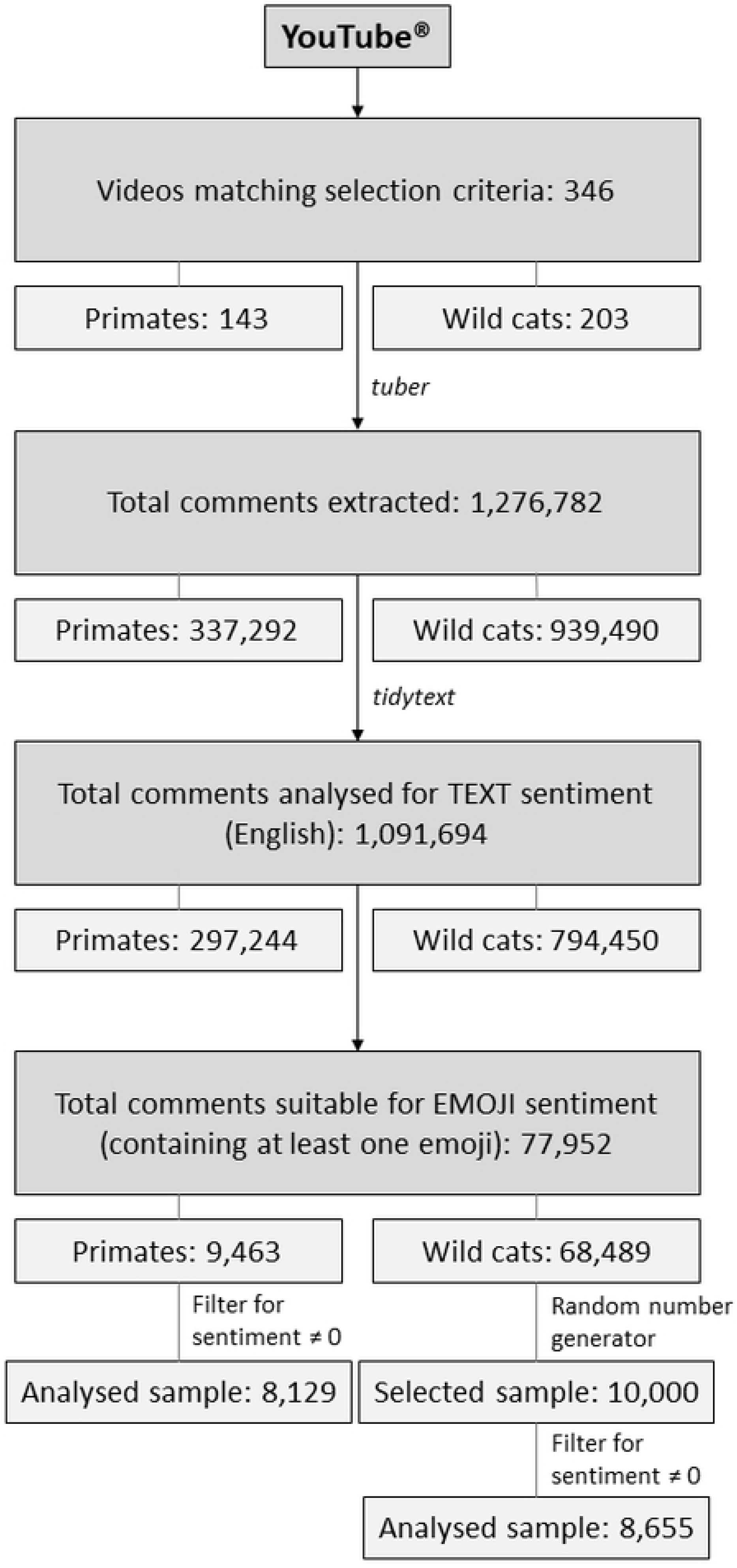
Methodological framework for comment extraction and analysis. Methodological framework for comment extraction and analysis across the 346 YouTube^®^ videos published from 2006 to 2019, sourced in accordance with pre-determined selection criteria. Only comments extracted written in English, or containing terms identified by the sentiment software employed, were analysed for text sentiment. Likewise, only comments containing at least one emoji were analysed for emoji sentiment. As wild cat emoji-containing comments were over represented within the dataset, a random sample of 10,000 comments was selected for emoji sentiment analysis. Filters were applied to ensure comments analysed for emoji sentiment contained at least one emoji and had an associated positive or negative sentiment score.

## Results

### Frequency analysis

In total, 346 videos were compiled for analysis with the majority (n = 203, 58.7%) categorised as exotic wild cat videos, based on the prominent species depicted and search terms through which they were sourced. The most popular species recorded was the tiger (*Panthera tigris*), making an appearance in 48.3% of all wild cat videos (Table 1).

**Table 1:** Species representation and associated conservation status.

| Species | Frequency | Percentage (%)<sup>1</sup> | CITES<sup>2</sup> | IUCN<sup>3</sup> Red List |
| --- | --- | --- | --- | --- |
|  |  |  | Appendix | Status |
| Wild cats<sup>4</sup> (a) |  |  |  |  |
| Tiger (*Panthera tigris*) | 98 | 48.3 | I | Endangered |
| Lion (*Panthera leo*) | 64 | 31.5 | II | Vulnerable |
| Cheetah (*Acinonyx jubatus*) | 53 | 26.1 | I | Vulnerable |
| Jaguar (*Panthera onca*) | 13 | 6.4 | I | Near Threatened |
| Leopard (*Panthera pardus*) | 13 | 6.4 | I | Vulnerable |
| Caracal (*Caracal caracal*) | 3 | 1.5 | I | Least Concern |
| **Primates<sup>5</sup> (b)** |  |  |  |  |
| Capuchin monkey (*Cebus apella*) | 42 | 29.4 | II | Least Concern |
| Macaque (*Macaca spp.*) | 28 | 19.6 | I | Endangered |
| Vervet monkey (*Chlorocebus pygerythrus*) | 8 | 5.6 | II | Least Concern |
| Chimpanzee (*Pan troglodytes*) | 7 | 4.9 | I | Endangered |
| Gibbon (*Nomascus*) | 6 | 4.2 | I | Critically Endangered |
| Pygmy slow loris (*Nycticebus pygmaeus*) | 7 | 4.9 | I | Vulnerable |
| Orangutan (*Pongo*) | 6 | 4.2 | I | Critically Endangered |
| White faced capuchin monkey (*Cebus albifrons*) | 6 | 4.2 | II | Least Concern |
| Green monkey (*Chlorocebus sabaeus*) | 5 | 3.5 | II | Least Concern |
| Spider monkey (*Ateles*) | 5 | 3.5 | I | Endangered |
| Pygmy marmoset (*Callithrix pygmaea*) | 4 | 2.8 | II | Least Concern |
| Gorilla (*Gorilla*) | 3 | 2.1 | I | Critically Endangered |
| Black and white ruffed lemur (*Varecia variegata*) | 3 | 2.1 | I | Critically Endangered |
| Squirrel monkey (*Saimiri*) | 2 | 1.4 | I | Vulnerable |

With regards to the wild cat videos, tigers attracted the greatest amount of comments, followed by lions (*Panthera leo*) then cheetahs (*Acinonyx jubatus*), closely reflecting species frequencies within the dataset (Table 1). Conversely, with regards to the primate videos, vervet monkeys (*Chlorocebus pygerythrus*) attracted the highest number of comments, followed by capuchin monkeys (*Cebus apella*) and macaques (*Macaca*).

The number of instances and percentage of corresponding YouTube^®^ videos associated with select search terms published between 2006 and 2019 wherein key exotic wild cat (a, n = 203) and primate (b, n = 143) species were identified. All videos required a minimum of one million views with the comments section enabled. A multiple response analysis was considered, wherein some videos featured multiple species. Instances wherein primate or exotic wild cat videos featured more than one associated species have been included. Species with a frequency equal to or less than one have been excluded. Primate compilation videos (n = 12) wherein species were not specified have also been excluded.

1. Percentage of wild cat (a, n = 203) or primate (b, n = 131, excluding ‘compilation’ videos) videos which featured a particular species
2. Convention on International Trade in Endangered Species of Wild Fauna and Flora (CITES) Appendix entries (2019);
3. International Union for Conservation of Nature (IUCN, 2019);
4. Other wild cat species featured include serval (*Leptailurus serval*), puma (*Puma concolor*);
5. Other primate species featured include baboon (*Papio spp.*), tarsier (*Tarsius spp.*), owl monkey (*Aotus spp.*)

Humans interaction was observed in 90.5% of all videos. The most popular non-human species interaction depicted was with domestic dogs which featured in 17.9% of all videos.

The predominant language detected within comments was English (49.4%). Of the comments from which emojis were extracted, emoji usage varied between the animal groups. Overall, 8.6% of wild cat comments contained at least one emoji, compared with only 3.2% of primate comments.

Species listed in CITES Appendices I and II were represented within the dataset, indicating imposed trade restrictions [5]. Likewise, species IUCN Red List conservation statuses ranged from ‘Least Concern’ to ‘Critically Endangered’.

#### Text sentiment analysis

The most popular sentiment terms identified in both wild cat and primate comments were ‘cute’, ‘like’ and ‘love’. ‘Like’ presented as the most frequently used term associated with wild cats, comprising 8.6% of all sentiment terms recognised, compared to ‘cute’ constituting 11.5% of all sentiment terms identified within primate comments.

### Emoji sentiment analysis

From the sample of 8,655 analysed wild cat emoji-containing comments (derived from 10,000 randomly selected comments), 87.1% (n = 7,540) were classified as positive based on a mean emoji sentiment score of between 0 and 1, compared with 89.8% (n = 7,299) of the 8,129 analysed primate emoji-containing comments.

### Comment analysis and conservation status

All mean emoji sentiment scores for wild cat (Fig 2a) and primate (Fig 2b) species IUCN Red List classifications remained above zero, suggesting positive public perception of these animals in selected YouTube^®^ videos despite severity of conservation status. Note that not all IUCN classifications were equally represented within the dataset.

**Fig 2.**
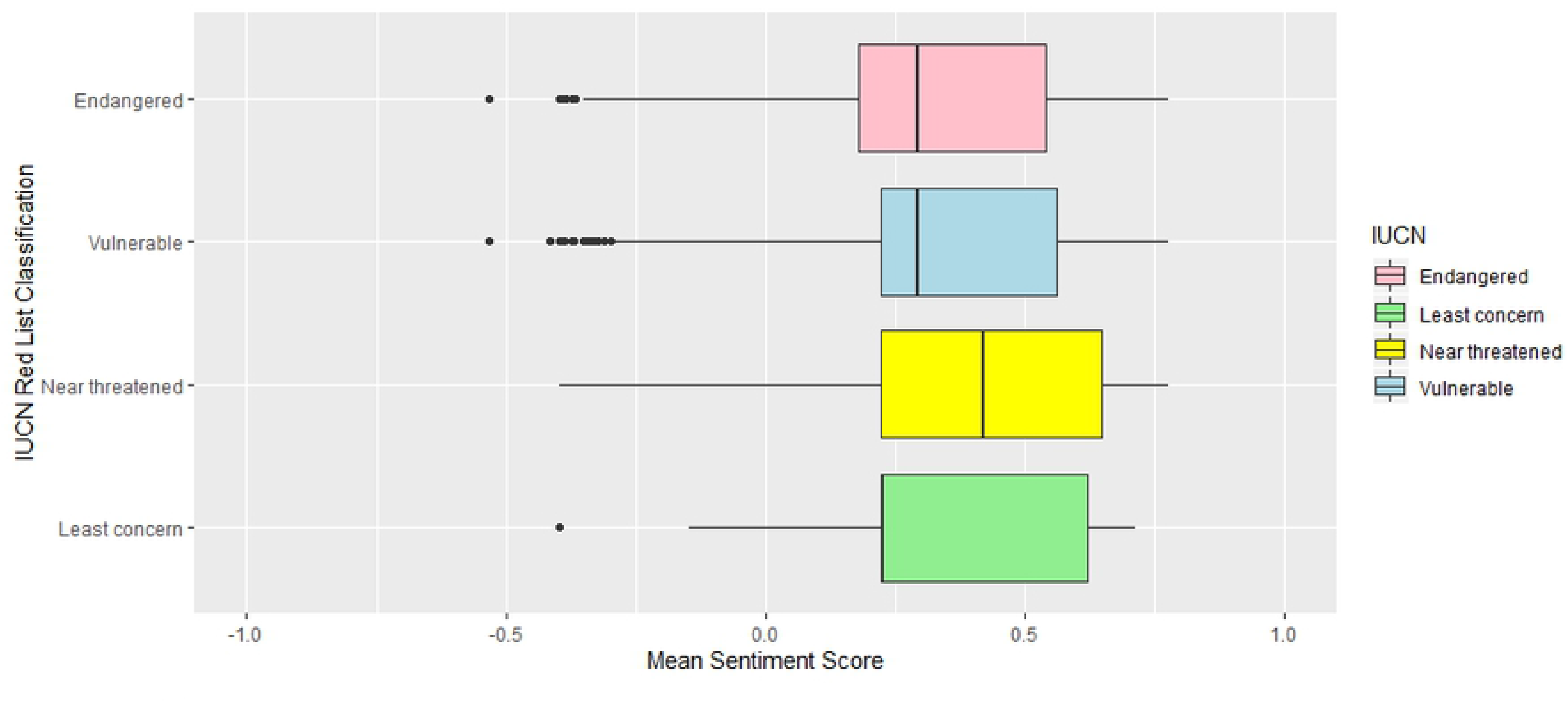
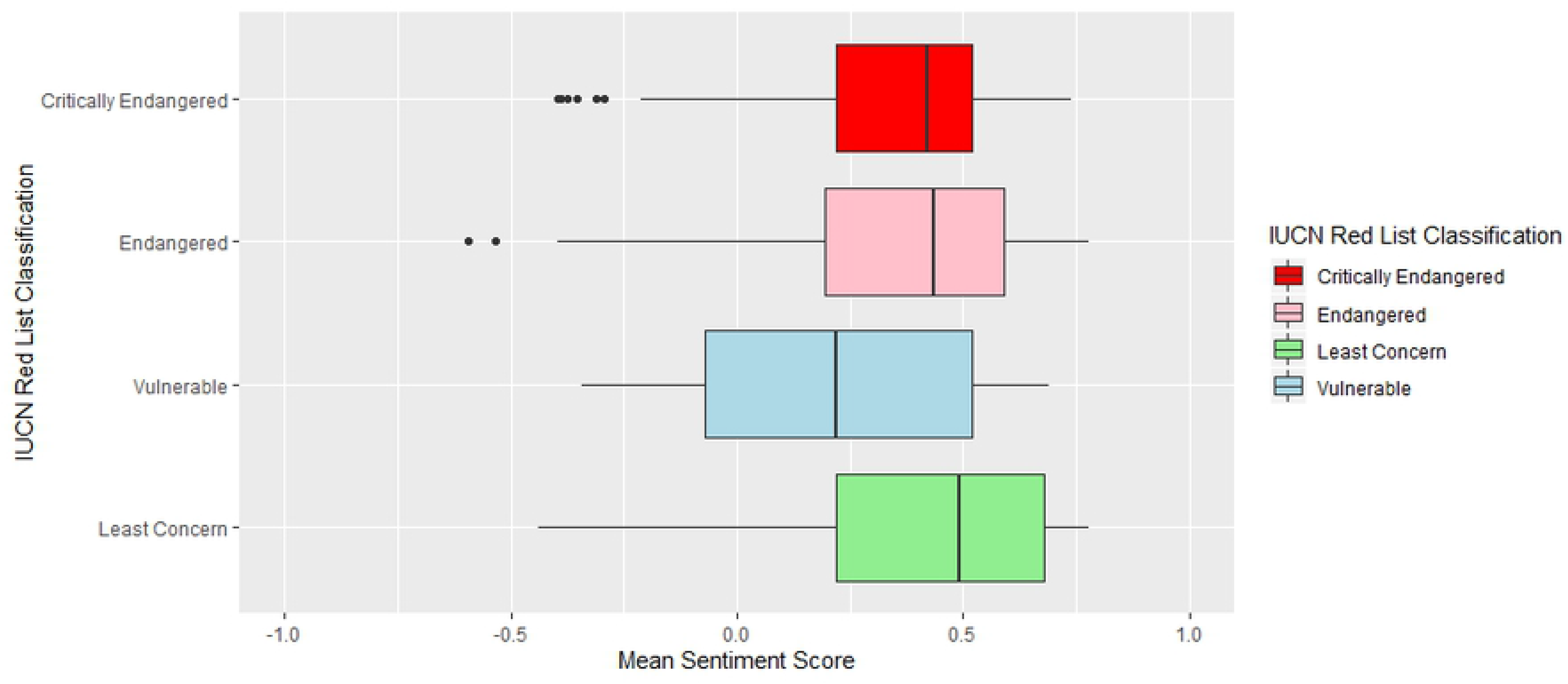
Emoji sentiment and IUCN Red List Classification. Mean emoji sentiment score across exotic wild cat (top graph, a; n = 8,655) and primate (bottom graph, b; n = 8,129) comments published between 2012 and 2019 associated with species conservation status as listed on the IUCN Red List (2019). Emoji usage within the comments was only present from 2012. Twelve primate compilation videos featuring four or more compiled videos featuring various species were not provided with an IUCN Red List classification due to ambiguity and as such have not been included. Mean sentiment score ranged between −1 (negative) and +1 (positive), where 0 is neutral, in accordance with the software utilised.

### Comparison of text and emoji sentiment over time

Mean sentiment scores remained above zero through time, indicating a positive sentiment and hence perception across both exotic wild cats and primates (Fig 3a and 3b).

**Fig 3.**
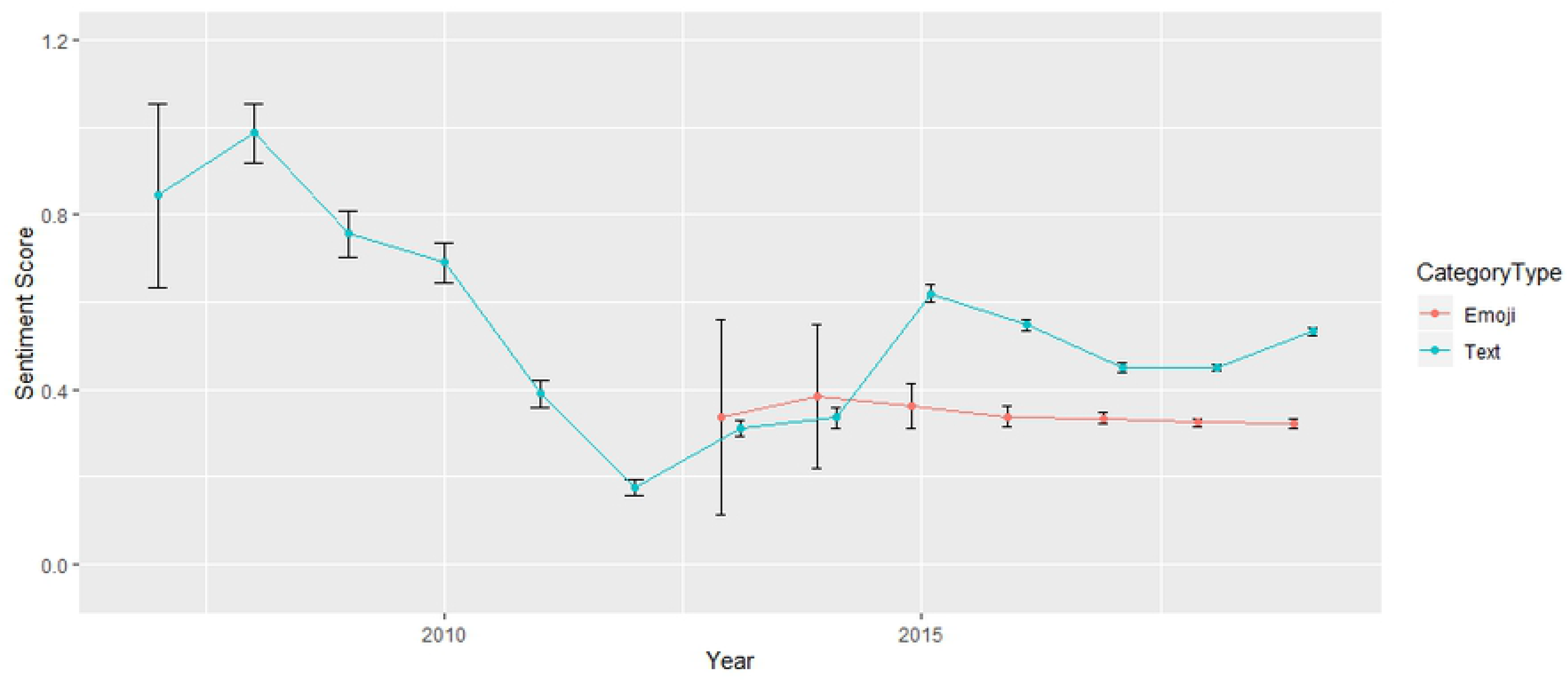
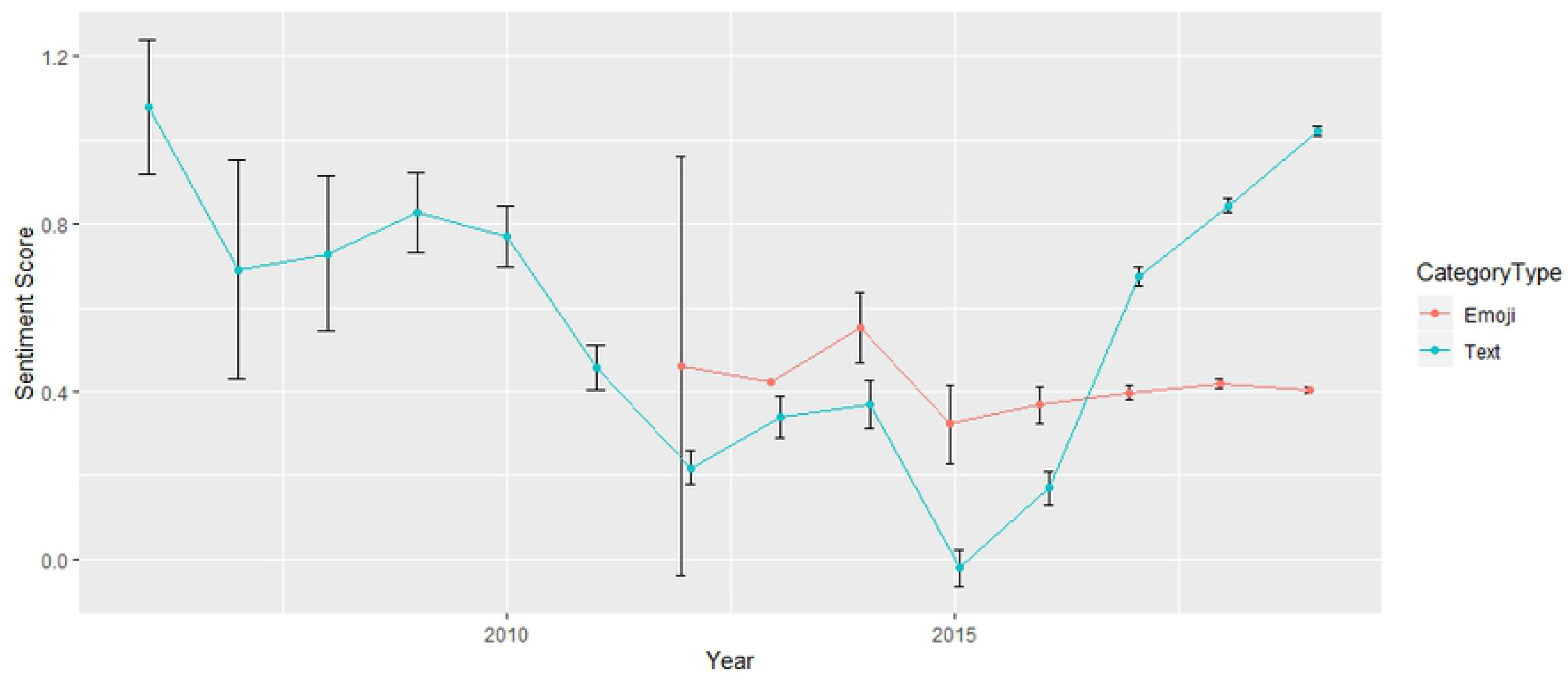
Text verses emoji sentiment through time. Mean (and 95% CI) text sentiment (blue) compared with mean (and 95% CI) emoji sentiment scores (red) in a random sample of wild cat comments (top graph, a; n = 8,655) and primate comments (bottom graph, b; n = 8,129) published from 2006 to 2019. Emojis were only present from 2012. Wild cat emoji sentiment for 2013 was only based on one observation (one emoji-containing comment associated with one video), hence no confidence intervals were applied for this year (3a). Mean primate emoji sentiment score was removed from 2012 as it was only based on two observations and was associated with an extreme confidence interval (3b).

Comparison of wild cat text and emoji sentiment displays similar trends, particularly during the period from 2012 to 2014, however there does not appear to be a strong association between them. Similarly, comparison of primate text and emoji sentiment over time displays similar trends. A negative mean sentiment was observed in 2015 for primates (Fig 3b), but not for wild cats (Fig 3a). Overall, both text and emoji mean sentiment scores remained consistently positive.

## Discussion

This study explored public perception of exotic species featured in popular YouTube^®^ videos and variation in perception throughout the life of a video. Potential association between exotic species in YouTube^®^ videos, the exotic pet trade and impact on species conservation will be discussed. Overall, both text and emoji sentiment analysis highlighted a predominately positive public response to content featured, despite species conservation status, which may subsequently influence trade in these species.

### Sentiment analysis

The prevailing frequency of positive terms, especially ‘cute’, ‘like’ and ‘love’, within video comments reflects the use of the ‘cute’ factor as a marketing tool for exotic pets [4,24]. However, limitations associated with the sentiment software were identified, including the inability to assess non-English words or to recognise word usage in alternate contexts, such as sarcasm [14,25]. To overcome these limitations, emoji usage was assessed, wherein the overwhelming majority of analysed comments (87.1% wild cats, 89.8% primates) indicated positive global public perception. Yet limitations associated with the ability to distinguish between alternate uses for emojis may have provided inaccuracies, thus further research may be required to validate its application in this type of research [25].

### Comment analysis and conservation status

Analysis of emoji usage demonstrated a varied spread of mean sentiment scores across IUCN Red List classifications (Fig 2). The absence of a significant association between sentiment score and severity of conservation status potentially highlights the public’s limited awareness or understanding of the associated welfare implications. It may be argued that portraying exotic animals interacting with humans, as seen in 90.5% of all videos, alludes to the supposed domestication of such animals and consequently endorses exotic pets [26,27]. Additional research supports the notion that displaying threatened exotic species in domestic settings commonly results in the misconception of their suitability as pets and affects perception of their wild populations status [26].

Interestingly, the most viewed video included in the study displayed ‘Endangered’ macaques performing for entertainment purposes (S2 Table). Despite their conservation status and evident animal welfare implications, the associated volume of views and overall positive sentiment score suggested this content was positively received by the audience. The evident lack of understanding mirrors findings presented by Stazaker and Mackinnon [28], where an assumed public ignorance of species conservation status may translate into creating demand for these animals.

The exploitation of wild animal populations has significantly contributed to welfare issues and conservation loss [2,24]. Due to the consistent positive public perception highlighted, it is suggested that the public are inadequately educated about origins of featured animals and the associated welfare consequences. Education is vital to ensure viewers not only identify the inappropriate depiction of exotic animals, but are also encouraged to report such content under YouTube^®^’s current guidelines [11].

### Impact on the exotic pet trade

Accessibility to the sale of exotic animals has been revolutionised through the internet, where content has supported the normalisation, desirability and acceptability of exotic species as pets [24]. Compared to primates, wild cats were the most popular group represented within the dataset. Tigers appeared in 48.3% of all wild cat videos in a wide range of domestic and captive situations despite their ‘Endangered’ conservation status [29,16], wherein their popularity on social media is supported by Spee et al. [27]. Exotic cats are heavily traded due to their popularity as pets, allure in captivity and demand for their parts, where the internet is significantly involved in facilitating the trade [30].

Although majority of the species featured were listed under CITES Appendix I, wherein trade is only permitted under exceptional circumstances, many have been reported in the live trade [5]. Harrington [31] reported popular wild cat and primate species traded and reported via CITES included lions, tigers, leopards, jaguars, servals, squirrel monkeys, marmosets, capuchins and tamarins, many of which were sourced from wild populations. These particular species were also featured in selected YouTube^®^ videos (Table 1). Trade routes for the importation and export of live animals were identified to include South America, North America, Africa, Asia and Europe, demonstrating the vast reach of the live exotic animal trade, wherein this distribution was reflected in the video origins included within this study (S2 Table) [31]. Regardless of CITES protection status and trade restrictions, there is still an extensive illegal trade present to supply the significant demand for exotic pets, much of which remains unmonitored [24]. Understanding how social media influences public perception and consumer demand is essential to enable implementation of effective strategies to monitor the modern trade and exploitation of threatened species.

### Comment analysis through time and promoting public awareness

Mean emoji sentiment scores consistently remained positive throughout time for both wild cat and primate comments (Fig 3). Interestingly, even with the significant decline in primate text sentiment in 2015, emoji sentiment followed a similar trend but continued to remain positive (Fig 3b). However, emojis are more likely to be used in association with expressing positive sentiment, which may bias the results [12]. Campaigns and initiatives pioneered by legitimate organisations, such as the “Tickling is TORTURE” campaign launched in 2015 which aimed to combat slow loris exploitation, have had an evident impact on the perception of exotic animals featured online [32]. The success of this strategy may even be reflected in the negative trend associated with primate videos in 2015 (Fig 3b), highlighting the impact of public education. However, following this decline sentiment scores began to increase, possibly suggesting the awareness encouraged by the campaign was either forgotten or not extrapolated to other exotic species. Fluctuating internet trends and ‘viral videos’ need to be closely monitored as they challenge the implementation of strategies to successfully oppose featured content [33].

### YouTube^®^ policies and content

Content posted on YouTube^®^ is loosely regulated, despite the provision of policies outlining expectations and limitations, wherein the site heavily relies on consumers to report (‘flag’) policy violations or illegal content [11]. In the case of animal abuse, reporting under the “violent or graphic content policies” guidelines (S1 Fig) is heavily reliant on an individuals’ ability to detect signs of distress or mistreatment, which may vary. As indicated by this research, the public cannot be relied upon to accurately identify inappropriate use of exotic animals, as much of the content featured was seemingly considered ‘acceptable’ despite underlying welfare and conservation concerns. As YouTube^®^ itself plays a crucial role in enabling public access to this content, they must also accept social responsibility for creating a culture wherein human engagement with threatened exotic wildlife has become acceptable.

Effective content monitoring can not only assist with gauging public perception to address the evident education gaps, but also help to identify new trends within the exotic pet industry and thus infer conservation and public health risks. YouTube^®^ policy guidelines should be updated to prohibit videos displaying interactions between humans and exotic wildlife, due to the perception of normalisation promoted. It is further recommended YouTube^®^ employ software to automatically detect key terms, such as species names, within video titles or descriptions and flag them for review, wherein videos in violation of newly established policies should be promptly removed. Alternatively, the development of an advertisement campaign presenting information surrounding exotic animal exploitation through the pet trade and the importance of species conservation could be automatically applied before all exotic animal videos to discourage support from viewers.

It is further suggested that the platform enables the provision of information to viewers to allow them to make a conscious decision when accessing content, similar to Instagram^®^’s ‘Wildlife Alert System’ designed to combat the use of exotic animal photo props [34]. However, in relation to Moorhouse et al. [35], it may be beneficial to focus information around the potential zoonotic risks and legal consequences of exotic pet ownership in addition to the conservation implications to better engage viewers. A clear icon could appear above each video which viewers could select, revealing current conservation and welfare information, as well as how to report videos in violation of newly established policy guidelines.

## Conclusions

Analysis of text and emoji usage within comments revealed a predominantly positive global public perception in response to the exploitation of exotic wild cats and primates on YouTube^®^. Although exotic species featured ranged in CITES listings and IUCN conservation status, the overall positive response generated appeared to indicate a lack of public awareness. In response, implementation of targeted YouTube^®^ policies, provision of information for users and accurate content regulation is highly encouraged to cease the illusion of the normalisation of threatened exotic animals as pets and prevent false legitimisation of the trade.

## Supporting information

**S1 Table. YouTube search terms.** Search terms investigated through the YouTube^®^ search engine to source videos for analysis. Only search terms which produced new videos selected for analysis are listed.

**S2 Table. Popular videos within the study.** The top ten most popular videos included within the study, according to view count at the time of data extraction, listed with primary species featured and additional video specifications.

**S1 Fig: YouTube^®^ policy guidelines.** Excerpts from YouTube^®^’s “violent or graphic content” policy guidelines available online. The policy sections featured are those pertaining to the use of animals within videos.

## References

1. Nijman V. An overview of international wildlife trade from Southeast Asia. Biodivers Conserv. 2010;19:1101–1114. doi: 10.1007/s10531-009-9758-4.

2. Bush ER, Baker SE, Macdonald DW. Global Trade in Exotic Pets 2006-2012. Biol. Conserv. 2014;28(3):663–676. doi: 10.1111/cobi.12240.

3. Baker SE, Cain R, van Kesteren F, Zommers ZA, D’Cruze N, Macdonald DW. Rough Trade: Animal Welfare in the Global Wildlife Trade. Bioscience. 2013;63(12):928–938. doi: 10.1525/bio.2013.63.12.6.

4. Siriwat P, Nijman V. Illegal pet trade on social media as an emerging impediment to the conservation of Asian otter species. J Asia-Pac Biodivers. 2018;11(4):469–475. doi: https://doi.org/10.1016/j.japb.2018.09.004.

5. CITES [Internet]. Appendices I, II and III; 2019 [cited 2019 July 4]. Available from: https://www.cites.org/eng/app/appendices.php.

6. Bergström A, Belfrage MJ. News in Social Media. Digital Journalism. 2018;6(5):583–598. doi: 10.1080/21670811.2018.1423625.

7. Kalogeropoulos A. Online News Video Consumption. Digital Journalism. 2018;6(5):651–665. doi: 10.1080/21670811.2017.1320197.

8. Khan ML. Social media engagement: What motivates user participation and consumption on YouTube? Comput. Hum. Behav. 2017;66:236–247. doi: http://dx.doi.org/10.1016/j.chb.2016.09.024.

9. Kumar S, Shah N. False Information on Web and Social Media: A Survey. arXiv:1804.08559 [Preprint]. 2018 [cited 2019 Aug 8]. Available from: https://arxiv.org/abs/1804.08559.

10. Chen Y, Chang C. Early prediction of the future popularity of uploaded videos. Expert Syst Appl. 2019;133:59–74. doi: https://doi.org/10.1016/j.eswa.2019.05.015.

11. YouTube [Internet]. Press; c2020 [cited 2019 Aug 8]. Available from: https://www.youtube.com/about/press/.

12. Prada M, Rodrigues DL, Garrido MV, Lopes D, Cavalheiro B, Gaspar R. Motives, frequency and attitudes toward emoji and emoticon use. Telemat. Inform. 2018;35(7):1925–1934. doi: https://doi.org/10.1016/j.tele.2018.06.005.

13. Ferrara HM. Sentiment Analysis. In: Gale Business Insights Handbook of Social Media Marketing. Cengage Learning; 2013. P. 325–337.

14. Novak PK, Smailović J, Sluban B, Mozetič I. Sentiment of Emojis. PLoS ONE. 2015;10(12). doi: 10.1371/journal.pone.0144296.

15. Nekaris KA, Campbell N, Coggins TG, Rode EJ, Nijman V. Tickled to Death: Analysing Public Perceptions of ‘Cute’ Videos of Threatened Species (Slow Lorises – *Nycticebus spp.*) on Web 2.0 Sites. PLoS ONE, 2013;8(7). doi: 10.1371/journal.pone.0069215.

16. IUCN [Internet]. The IUCN Red List of Threatened Species, Version 2019-3; 2019 [cited 2019 July 10]. Available from: http://www.iucnredlist.org.

17. RStudio Team. RStudio: Integrated Development for R. Version 3.6.1 [software]. 2015; Boston, MA. Available from: http://www.rstudio.com/.

18. Sood G. tuber: Access YouTube from R. R package version 0.9.8 [software]. 2019.

19. Hornik K, Mair P, Rauch J, Geiger W, Buchta C, Feinerer I. The textcat Package for n-Gram Based Text Categorization in R. J. Stat. Softw. 2013;52(6):1–17. doi: 10.18637/jss.v052.i06.

20. Silge J, Robinson D. tidytext: Text Mining and Analysis Using Tidy Data Principles in R. JOSS. 2016;1(3). doi: 10.21105/joss.00037.

21. Uryupina O, Plank B, Severyn A, Rotondi A, Moschitti A. SenTube: A Corpus for Sentiment Analysis on YouTube Social Media. Proceedings of the Ninth International Conference on Language Resources and Evaluation; 2014 May; Reykjavik, Iceland.

22. Mullen LA. Fast, Consistent Tokenization of Natural Language Text. J. Open Source Softw. 2018;3(23):655. doi: 10.21105/joss.00655.

23. Peterka-Bonetta J. Emoji Analysis in R. 2017 [cited 2019 May 2]. Available from: http://opiateforthemass.es/articles/emoji-analysis/.

24. World Animal Protection. Wild at heart: The cruelty of the exotic pet trade. 2019 [cited 2019 Oct 10]. Available from: https://www.worldanimalprotection.org.au/sites/default/files/media/au_files/wild-at-heart-report-2019.pdf.

25. Tian Y, Galery T, Dulcinati G, Molimpakis E, Sun C. Facebook Sentiment: Reactions and Emojis. Proceedings of the Fifth International Workshop on Natural Language Processing for Social Media; 2017 April 3; Valencia, Spain.

26. Kitson H, Nekaris K. Instagram-fuelled illegal slow loris trade uncovered in Marmaris, Turkey. Oryx. 2017;51(3):394–394. doi: 10.1017/S0030605317000680.

27. Spee LB, Hazel SJ, Dal Grande E, Boardman WSJ, Chaber AL. Endangered Exotic Pets on Social Media in the Middle East: Presence and Impact. Animals. 2019;9(8):480. doi: 10.3390/ani9080480.

28. Stazaker K, Mackinnon J. Visitor Perceptions of Captive, Endangered Barbary Macaques (*Macaca sylvanus*) Used as Photo Props in Jemaa El Fna Square, Marrakech, Morocco. Anthrozoos. 2018;31(6):761–776. doi: https://doi.org/10.1080/08927936.2018.1529360.

29. Goodrich J, Lynam A, Miquelle D, Wibisono H, Kawanishi K, Pattanavibool A, et al. Panthera tigris. The IUCN Red List of Threatened Species; 2015 [cited 2019 Oct 27]. Available from: https://www.iucnredlist.org/species/15955/50659951. e.T15955A50659951.

30. Nijman V, Morcatty T, Smith JH, Atoussi S, Shepherd CR, Siriwat P, Nekaris A, Bergin D. Illegal wildlife trade – surveying open animal markets and online platforms to understand the poaching of wild cats. Biodiversity. 2019;20(1):58–61. doi: 10.1080/14888386.2019.1568915.

31. Harrington LA. International commercial trade in live carnivores and primates 2006–2012: response to Bush et al. 2014. Biol. Conserv. 2015;29(1):293–296. doi: 10.1111/cobi.12448.

32. International Animal Rescue [Internet]. Slow Loris Rescue; c2019 [cited 2019 Sept 27]. Available from: https://www.internationalanimalrescue.org/slow-loris-sanctuary.

33. Broxton T, Interian Y, Vaver J, Wattenhofer M. Catching a viral video. J Intell Inf Syst. 2013;40(2):241–259. doi: 10.1007/s10844-011-0191-2.

34. Mills G. Social media giant takes action on wildlife ‘selfies’. Vet Rec. 2017;181(25):672–673.

35. Moorhouse TP, Balaskas M, D’Cruze NC, Macdonald DW. Information Could Reduce Consumer Demand for Exotic Pets. Conserv Lett. 2016;10(3):337–339. doi: 10.1111/conl.12270.

